# Bioactive produced by *Enterococcus Faecalis* targets IL-23 signalling and protects against colitis and joint disease

**DOI:** 10.64898/2026.07.21.739719

**Authors:** Rabina Giri, Anne-Sophie Bergot, Páraic Ó Cuív, Mark Morrison, Ranjeny Thomas, Jakob Begun

## Abstract

The IL-23/Th17 axis is a central driver of intestinal and spondyloarthritic inflammation, yet upstream regulatory mechanisms linking microbial signals to IL-23 production remain incompletely defined. NF-κB signalling, particularly via the c-Rel subunit, is a critical transcriptional regulator of IL23A (p19), positioning c-Rel as a nodal checkpoint in mucosal inflammation.

Here, we demonstrate that cell-free supernatant derived from *Enterococcus faecalis* AHG0090 (AHG0090-CS) suppresses c-Rel–dependent IL-23 signalling and attenuates inflammatory pathology across murine models of gut and joint disease. In the ZAP-70 mutant SKG model of spondyloarthritis and ileitis, AHG0090-CS significantly reduced weight loss, joint scores, and histological gut inflammation following curdlan challenge. In the Winnie model of spontaneous colitis, treatment similarly diminished inflammatory cytokine production.

Mechanistically, AHG0090-CS reduced IL-23p19 mRNA and protein expression in intestinal tissue and lamina propria myeloid cells, accompanied by decreased nuclear c-Rel intensity. Suppression extended to downstream IL-23–associated cytokines including IL-17A, GM-CSF, MCP-1 and IL-6. In human peripheral blood mononuclear cells and macrophages, AHG0090-CS attenuated LPS-induced IL-23 and pro-inflammatory cytokine production, supporting translational relevance.

Collectively, these findings identify microbial modulation of c-Rel–dependent IL-23 signalling as a tractable mechanism to restrain gut–joint inflammation and highlight targeting upstream NF-κB pathways as a therapeutic strategy in IL-23–driven immune-mediated disease.

## Introduction

Intestinal immune responses are shaped by the microorganisms harboured in the mammalian gastrointestinal tract^1^. To maintain homeostasis, the intestinal immune system has co-evolved with the gut microbiota to balance tolerance to commensal organisms with protective immunity against pathogens^2^. Disruption of this equilibrium contributes to chronic immune-mediated diseases, including inflammatory bowel disease (IBD), such as ulcerative colitis and Crohn’s disease^3^, as well as inflammatory arthritis, including rheumatoid arthritis and spondyloarthritides (SpA) such as ankylosing spondylitis and reactive arthritis. These disorders are characterised by impaired epithelial barrier function, heightened pro-inflammatory cytokine signalling, and defects in immunoregulatory pathways^4–6^.

There is substantial mechanistic overlap between IBD and inflammatory arthritis, evidenced by shared genetic susceptibility loci and responsiveness to overlapping therapies, including tumour necrosis factor (TNF) inhibition^7^. IL-23 and the associated Th17 driven immune responses are a dominant inflammatory axis in both conditions. Genome-wide association studies have identified variants in *IL23R*, encoding the receptor for IL-23, associated with susceptibility to Crohn’s disease and ulcerative colitis^8^, as well as other immune-mediated diseases including psoriasis and psoriatic arthritis^9^, positioning IL-23 as a central inflammatory pathway across gut and joint disease. At a mechanistic level, IL-23 production is tightly regulated by NF-κB signalling, with the c-Rel subunit directly driving transcription of the IL23A (p19) gene in both murine and human myeloid cells, thereby linking upstream inflammatory signalling to downstream Th17 responses^10^.

Experimental and clinical studies further support a role for IL-23 in intestinal and joint inflammation. In murine models, spontaneous colitis in IL-10-deficient mice and T-cell transfer models is attenuated in the absence of IL-23p19, but not IL-12p40^11, 12^, while recombinant IL-23 accelerates disease and anti-IL-23p19 treatment ameliorates colitis. In the ZAP-70 mutant SKG mouse model, which develops SpA-like joint disease and Crohn’s-like ileitis following exposure to microbial β-1,3-glucan (curdlan), IL-23p19 blockade reduces both ileal and joint inflammation^13^. In humans, elevated IL-17 and IL-23 levels are detected in the serum and synovial tissue of patients with rheumatoid arthritis and SpA^14^ and are largely absent from healthy joints^15^, consistent with increased expression of IL-12 and IL-23 in inflamed intestinal mucosa from IBD patients.

In parallel with shared immune signalling pathways, growing epidemiological and translational evidence implicates the gut microbiome in both intestinal and joint disease^16^. Reduced microbial diversity and depletion of Baciliota, including *Faecalibacterium* and *Roseburia* species, are observed in IBD, SpA^4^, and rheumatoid arthritis compared with healthy controls^17, 18^. Furthermore, subsets of IBD patients are at increased risk of developing arthritis following gastrointestinal infections, with specific bacterial strains implicated in cartilage degradation^19, 20^. Faecal microbiota transplantation–induced remission in ulcerative colitis provides direct evidence for a causal role of the gut microbiome in disease^21, 22^. Nonetheless, microbiome compositional analyses fail to define the functional microbial products responsible for regulating inflammatory pathways *in vivo*.

Microbiome-derived bioactive molecules, including small molecules and peptides, represent an important mechanism through which gut bacteria communicate with the host immune system^23^. While the immune system senses pro-inflammatory microbial products such as pathogen-associated molecular patterns^2, 24^, several microbial-derived factors exert anti-inflammatory effects. These include the microbial anti-inflammatory molecule (MAM) from *Faecalibacterium prausnitzii*, which suppresses NF-κB signalling^25, 26^ and bioactive extracts from *Lactobacillus sakei* that ameliorate inflammation in experimental psoriasis^27^. Together, these findings suggest that bacterially derived bioactives may represent a tractable means of modulating upstream inflammatory pathways relevant to immune-mediated disease.

We previously isolated *Enterococcus faecalis* AHG0090, a commensal strain capable of producing a secreted peptide-like bioactive that suppresses NF-κB activation and pro-inflammatory cytokine production *in vitro*^28^. However, whether this bioactive can modulate inflammation *in vivo* and across distinct disease contexts remained unknown. In this study, we investigated the immunomodulatory effects of cell-free culture supernatants from *E. faecalis* AHG0090 in two preclinical models of immune-mediated disease: spontaneous colitis in Winnie mice and curdlan-induced Crohn’s-like ileitis with spondyloarthritis in SKG mice. We demonstrate that bioactive production is culture-condition dependent and that administration of cell-free supernatant, but not live bacteria, attenuates intestinal and joint inflammation. Notably, this bioactive reduces IL-23 production by macrophages across both models and in IBD patient-derived explants, implicating modulation of the IL-23-c-Rel NF- κB axis as a shared inflammatory pathway relevant to gut and joint disease.

## Results

### Cell-free supernatant, but not live *E. faecalis* AHG0090, attenuates ileal and joint inflammation in the SKG mouse model

We first investigated whether *E. faecalis* AHG0090 could modulate inflammation *in vivo* using the curdlan-induced SKG mouse model, which develops Crohn’s-like ileitis and spondyloarthritis in an IL-23 and microbiota-dependent manner. To assess whether live bacterial administration was sufficient to confer protection, SKG mice with an intact gut microbiota were gavaged weekly with live *E. faecalis* AHG0090 prior to curdlan challenge. Live bacterial administration did not improve body weight loss, visual joint scores, or histological inflammation in the ileum or joints following curdlan exposure (Fig. S1A–F), indicating that delivery of intact bacteria alone was insufficient to ameliorate disease in this model.

Given that microbial immunomodulatory effects can depend on environmental context and nutrient availability, we examined whether production of bioactive factors by *E. faecalis* AHG0090 was influenced by bacterial growth conditions. *E. faecalis* AHG0090 was cultured in multiple media prior to preparation of cell-free culture supernatants (CS), and immunomodulatory activity was assessed using an NF-κB reporter assay. CS derived from bacteria grown in brain heart infusion (BHI) or M2GSC media significantly suppressed TNF-induced NF-κB activation, whereas CS generated from bacteria grown in MRS media lacked inhibitory activity (Fig. S1G). These data demonstrate that production of the immunomodulatory bioactive by *E. faecalis* AHG0090 is media dependent and identify BHI-derived CS as a robust source of bioactive activity for in vivo studies.

We therefore next assessed whether BHI-derived CS from *E. faecalis* AHG0090 could suppress inflammation in the SKG model following curdlan challenge. In contrast to live bacterial administration, treatment with bi-weekly intra-peritoneal AHG0090 CS injection after curdlan challenge significantly reduced resulting disease severity. In CS-treated mice body weight loss and visual arthritis scores were significantly lower than in untreated and BHI-treated curdlan-challenged SKG controls (Fig. 1A,B). Histological analysis of H&E sections revealed reduced inflammatory infiltrates, preservation of villus–crypt architecture in the ileum, and decreased synovial inflammation in ankle joints of CS-treated compared to control curdlan-challenged SKG mice (Fig. 1C,D). Consistent with these improvements, ileal expression of *Il23p19* and the ER stress marker *sXbp1* was significantly reduced in CS-treated compared to control mice (Fig. 1E). Together, these findings demonstrate that CS prepared from *E. faecalis* AHG0090 under appropriate growth conditions is sufficient to attenuate both intestinal and joint inflammation in curdlan-challenged SKG mice.

**Figure 1.**
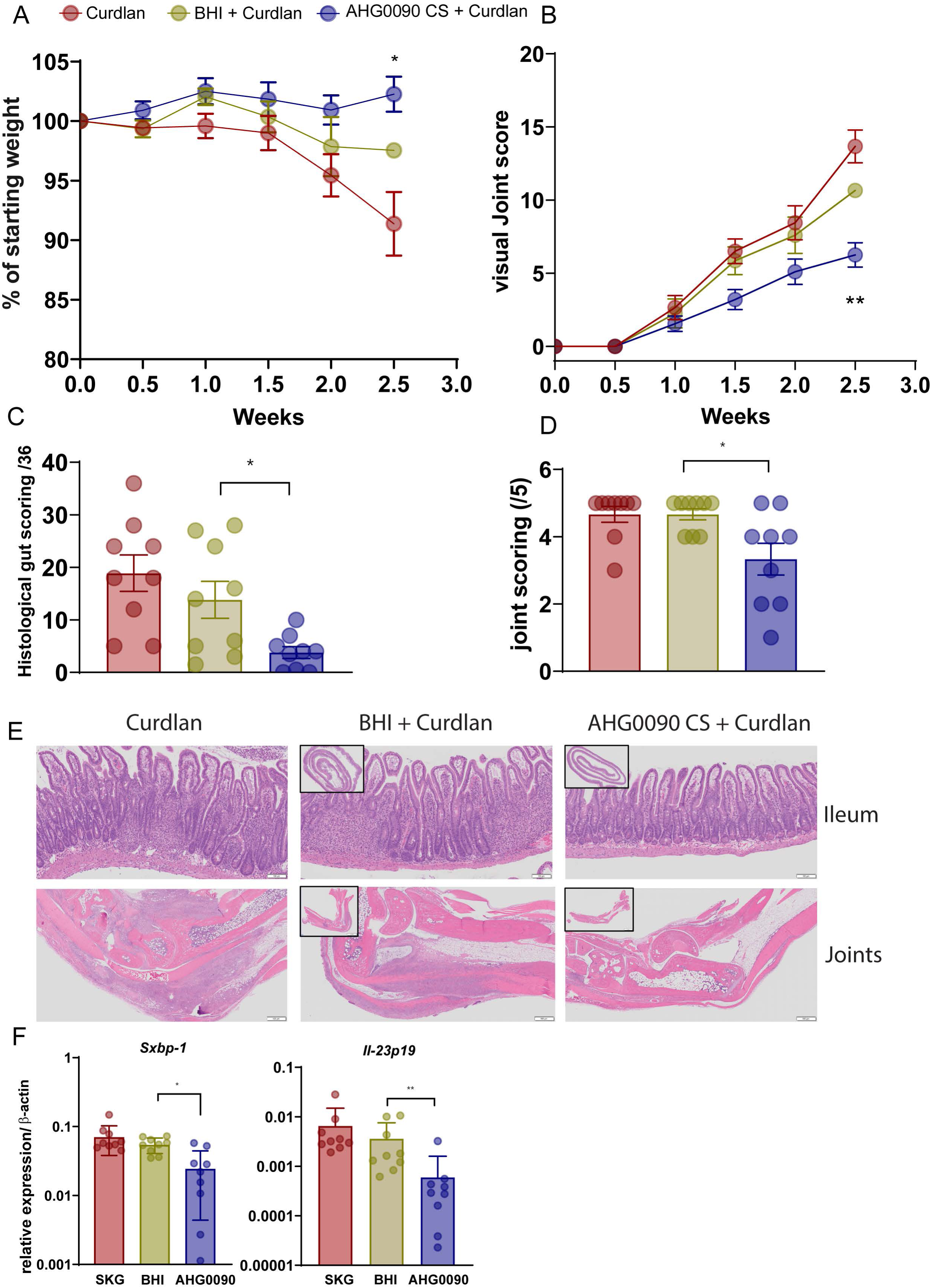
Culture supernatants from *E. faecalis* AHG0090 attenuate intestinal and joint inflammation in the SKG mouse model. SKG mice were challenged with curdlan on day -1 and treated with cell-free culture supernatant (CS) from *E. faecalis* AHG0090 or BHI control bi-weekly (at weeks 0.5, 1, 1.5, 2 & 2.5). (A) Changes in body weight over the course of disease. (B) Visual arthritis scores assessed throughout the experimental period. (C–D) Blinded histological scoring of inflammation in the ileum and ankle joints, with representative haematoxylin and eosin (H&E) stained sections shown for each tissue. (E) Ileal mRNA expression of inflammatory cytokines and the endoplasmic reticulum stress marker *sXbp1*. Each dot represents an individual mouse. Data are pooled from two independent experiments. Statistical significance was determined by one-way ANOVA with Dunnett’s multiple comparison test. *P* < 0.05, P < 0.01, *P* < 0.001, P < 0.0001.

### CS from *E. faecalis* AHG0090 ameliorates spontaneous colitis in Winnie mice

To determine whether the anti-inflammatory effects of *E. faecalis* AHG0090 CS extended to a model of intestinal colitis, we next evaluated its efficacy in Winnie mice, which develop spontaneous colitis driven by epithelial barrier dysfunction and endoplasmic reticulum stress due to a point mutation in the *Muc2* gene. Diarrhoea scores (Fig. 2A) and colon weight-to-length ratios were significantly reduced in Winnie mice treated intrarectally with *E. faecalis* AHG0090 CS compared with BHI-treated controls (Fig. 2B). Histological analyses demonstrated reduced immune cell infiltration, preservation of goblet cells, and improved crypt architecture in CS-treated compared to control Winnie mice (Fig. 2C). In parallel, colonic expression of inflammatory chemokines (*Mip2, Cxcl10), Il23p19*, and the ER stress marker *sXbp1* was significantly reduced following CS treatment (Fig. 2D). These results indicate that *E. faecalis* AHG0090-derived bioactives suppress inflammation in both induced and spontaneous models of intestinal disease.

**Figure 2.**
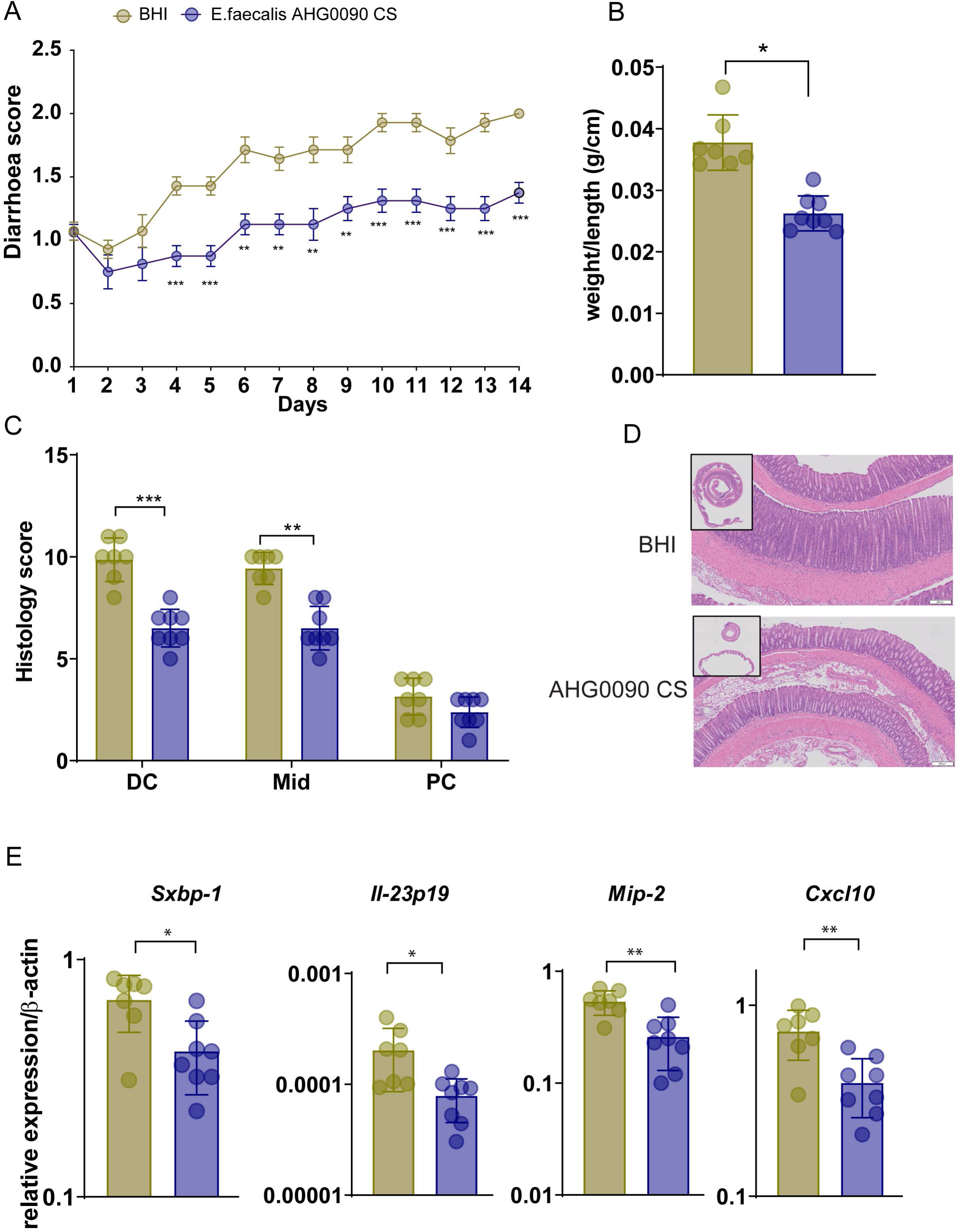
Cell-free culture supernatant from *E. faecalis* AHG0090 ameliorates spontaneous colitis in Winnie mice. Winnie mice were treated intrarectally with cell-free culture supernatant (CS) from *E. faecalis* AHG0090 or BHI control for 14 days. (A) Diarrhoea scores assessed during the treatment period (B) Colon weight-to-length ratio at experimental endpoint (C) Blinded histological scoring of colonic inflammation (D) Representative haematoxylin and eosin (H&E) stained sections of distal colon showing epithelial architecture and inflammatory infiltrates (E) Colonic mRNA expression of inflammatory chemokines (Mip2, Cxcl10), Il23p19, and the endoplasmic reticulum stress marker sXbp1. Each dot represents an individual mouse. Data are pooled from two independent experiments. Statistical significance was determined by one-way ANOVA with Dunnett’s multiple comparison test. P < 0.05, P < 0.01, P < 0.001, P < 0.0001.

### *E. faecalis* AHG0090 selectively suppresses IL-23 production by macrophages in murine inflammatory tissues

Given the reduction in IL-23 in both disease models, we sought the cellular sources targeted by AHG0090 CS. Treatment of colonic explants from Winnie mice and ileal explants from curdlan-treated SKG mice with *E. faecalis* AHG0090 CS significantly reduced secretion of IL-23, IFN-γ, MCP-1, and GM-CSF (Fig. 3A,B). At the transcriptional level, reductions in *Il23p19* and *sXbp1* expression were observed across all tissues (Fig. 3C,D).

**Figure 3.**
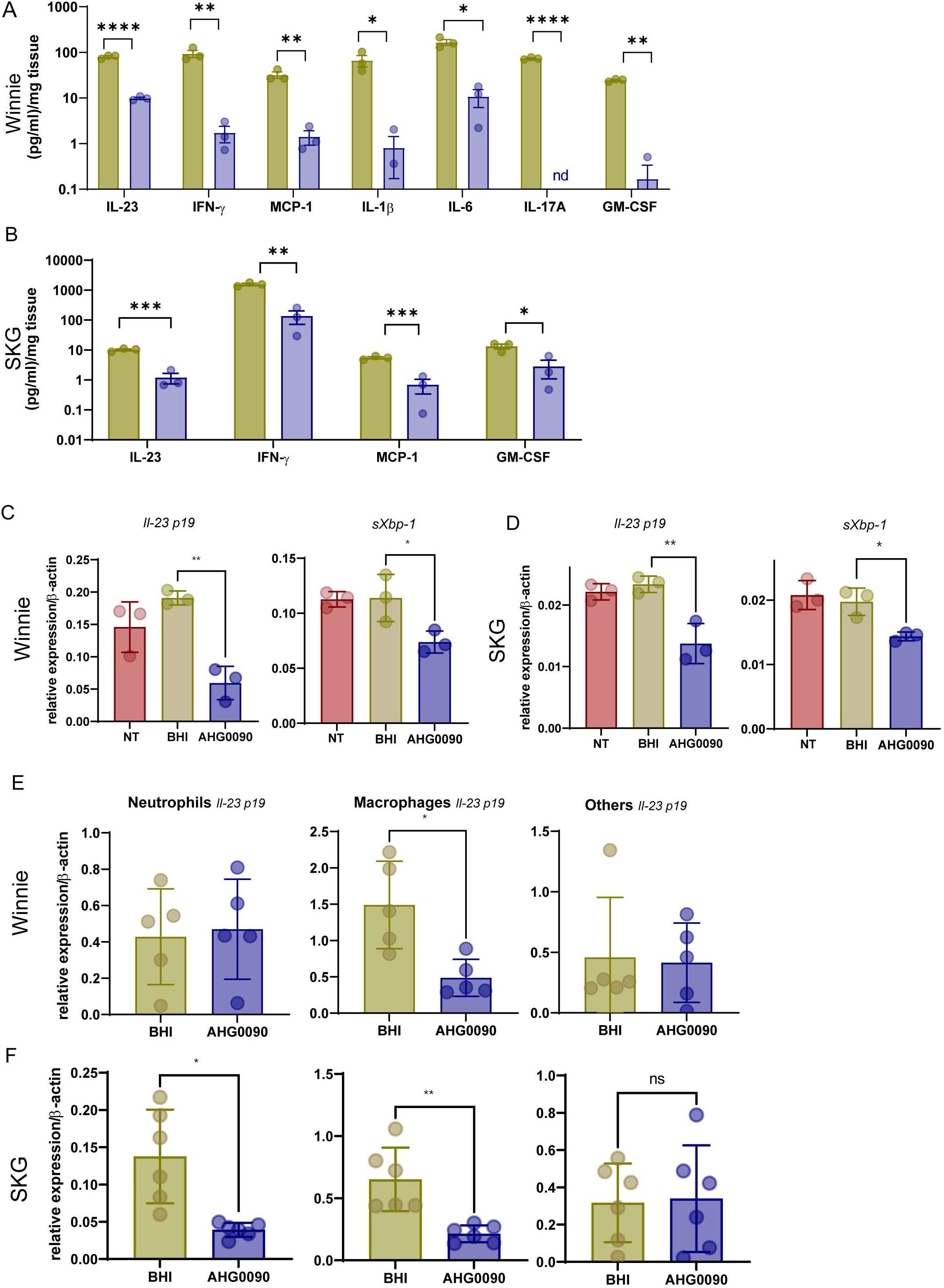
*E. faecalis* AHG0090 bioactive suppresses IL-23 production by macrophages in murine inflammatory tissues. A) Colonic explants from 12 weeks old Winnie mice and ileal explants from curdlan treated SKG mice were treated with either media only, 10% v/v BHI media or CS from *E. faecalis* AHG0090 for 24 hours *ex vivo.* Cytokines and chemokines secretions were measured. Cytokine/chemokine concentration was normalised to weight (mg) of tissue. C-D) Explants were treated for 6 hours and mRNA expression of Il-23p19 and sXBP-1 were determined using qPCR E) Il-23p19 mRNA levels from sorted populations from Winnie lamina propria and exudate from SKG mice. Each circle represents individual mouse.

Given the previously described role of IL-23 in both the SKG and Winnie mouse models, and the significant suppression of *Il23p19* expression, we explored the effects of *E. faecalis* AHG0090 on this cytokine. As IL-23 is predominantly produced by myeloid cells, we next examined whether AHG0090 CS selectively modulated IL-23 production within this compartment *in vivo*. In Winnie mice, treated with CS for 1 hour, lamina propria immune cell composition was unchanged (Fig. S2A–C); however, *Il23p19* expression was reduced in macrophages but not in neutrophils or non-myeloid cells (Fig. 3E). Similarly, in SKG mice treated for 24 hours with CS, we observed reduced *Il23a* expression in peritoneal exudate macrophages and neutrophils (Fig. S3A-C) collected 6 hours after curdlan challenge(Fig. 3F). These changes were independent of ER stress modulation (*sXbp1* (Fig. S3D)). These findings identify macrophages, as a key cellular target of *E. faecalis* AHG0090 bioactive activity in the colon and peritoneal macrophages and neutrophils as acute cellular targets of AHG0090 after curdlan-challenge *in vivo*.

### *E. faecalis* AHG0090 CS suppresses IL-23 production in IBD patient-derived tissues and monocyte-derived macrophages

To assess translational relevance, we next examined the effects of *E. faecalis* AHG0090 CS on human IBD samples. Secretion of IL-23 and IL-6, but not IL-8 were significantly reduced in ileal and colonic explants from patients with Crohn’s disease and ulcerative colitis treated *ex vivo* with AHG0090 CS compared with BHI (Fig. 4A) (Fig. S4A,B), with inter-patient variability observed. In peripheral blood mononuclear cells, *E. faecalis* AHG0090 CS reduced LPS-induced IL-6 secretion (Fig. S4C). In monocyte-derived macrophages, *E. faecalis* AHG0090 CS significantly suppressed LPS-induced IL-23 and IL-6 production (Fig. 4B). These data demonstrate that *E. faecalis* AHG0090 CS modulates IL-23 and IL-6 in human immune and tissue contexts relevant to IBD.

**Figure 4.**
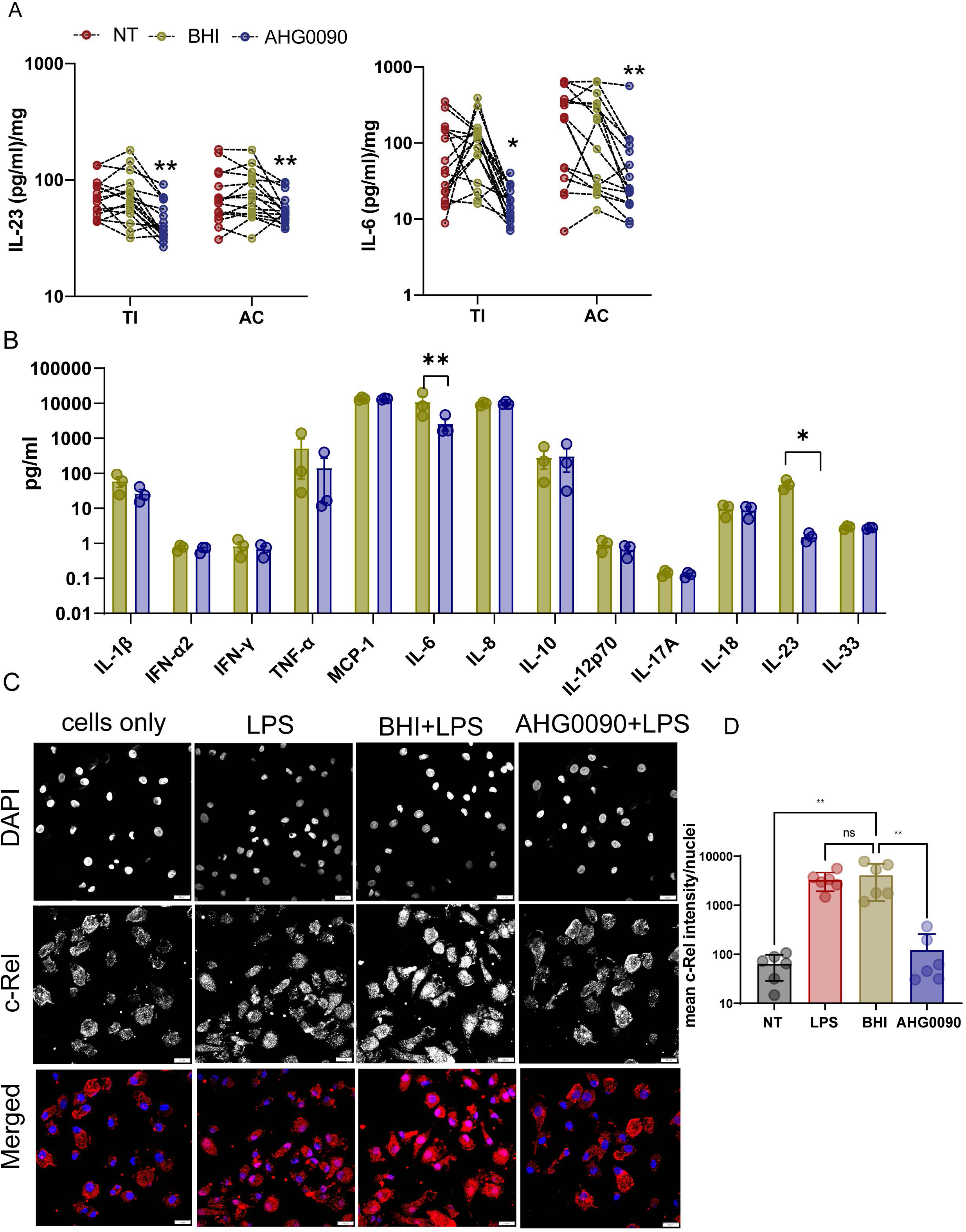
*E. faecalis* CS reduces IL-23 secretion in macrophages by inhibiting c-Rel translocation. **A)** Ileal and colonic biopsies from IBD subjects were treated with 10% v/v BHI or CS from *E. faecalis* AHG00090 for 24 hours and IL-6 and IL-23 were measured using ELISA. Each circle represents an individual subject. B) Monocyte derived macrophages from 3 non-IBD subjects were stimulated with LPS +/- CS from BHI or AHG0090 for 24 hours. Cytokines/Chemokines were measured using Legendplex Human Inflammation kit. C) Monocytes derived macrophages were treated with LPS +/- BHI or CS from AHG0090 for 30mins. Cells were fixed, permeabilized and stained with c-rel Antibody. Cells were visualised using Olympus FV3000 at 60x. Blue represents nuclei (DAPI) and red represents C-rel staining. Nuclear c-Rel staining was quantified using Image J. 2 representative images per subject (n=3) were quantified and > 10 nuclei per field were quantified.

### *E. faecalis* AHG0090 bioactive inhibits NF-κB c-Rel nuclear translocation in macrophages

IL-23p19 transcription is regulated downstream of toll-like receptor signalling and depends on NF-κB activity, particularly the c-Rel subunit, in myeloid cells. Given the consistent suppression of macrophage-derived IL-23 across murine models and human samples, and our prior observation that AHG0090 suppresses NF-κB activation *in vitro*, we hypothesised that the secreted *E.faecalis* AHG0090 bioactive interferes with NF-κB-dependent transcriptional regulation of *Il23p19*. Consistent with this hypothesis, *E. faecalis* AHG0090 CS markedly inhibited LPS-induced nuclear translocation of c-Rel in macrophages compared with BHI control treatment (Fig. 4C). These findings provide a mechanistic basis for reduced IL-23 production and implicate modulation of the IL-23–c-Rel- NF-κB axis as a pathway through which *E. faecalis* AHG0090-derived bioactive attenuate intestinal and joint inflammation.

## Discussion

In this study, we focused on *E. faecalis* AHG0090, a commensal strain that suppresses NF-κB activation *in vitro*^29^. We demonstrate that cell-free culture supernatants from *E. faecalis* AHG0090 attenuate inflammation in two mechanistically distinct preclinical models - spontaneous colitis in Winnie mice and curdlan-induced Crohn’s-like ileitis with spondyloarthritis in SKG mice – and in human tissue explants from IBD patients. These findings provide functional evidence that bioactive molecules derived from a prevalent human gut bacterium can modulate inflammation across immune-mediated diseases affecting both the gut and joints. Prior studies have reported conflicting roles for *E. faecalis* in colitis, with evidence for both pathogenic and protective effects depending on strain and context^30, 31^. Our data help reconcile these discrepancies by demonstrating that the production of functional bioactives, rather than bacterial presence alone, determines immunomodulatory potential. Supporting this, a recent study identified 12 genome-sequenced *Enterococcus* representative isolates of the IBD-associated and control clades (6 each) and found that control clade isolates induced significantly greater cell cytotoxicity than isolates from the IBD-associated clade, highlighting the importance of pairing microbial composition with its function and host transcriptional response^32^. In contrast to IBD, there have been limited studies on the role of *E. faecalis* in pathogenesis of arthritis, although a lower abundance of *E. faecalis* has been reported in stool from RA patients compared to healthy controls^33^.

Our *in vitro* and *ex vivo* data also suggests that the functional capacity of bacteria is dependent on their growth environment. While live *E. faecalis* AHG0090 failed to ameliorate disease in the SKG model, administration of cell-free supernatant robustly suppressed intestinal and joint inflammation. We further show that bioactive production is strongly dependent on bacterial growth conditions, consistent with prior observations using other immunomodulatory strains^34^. These data highlight a major limitation of live biotherapeutic approaches: microbial function is highly context-dependent and may not be reliably recapitulated *in vivo*. In contrast, bioactive-based strategies bypass challenges associated with engraftment, microbial competition, and nutrient availability.

Mechanistically, suppression of inflammation across both murine models and in human IBD tissues was accompanied by consistent reduction of IL-23, a cytokine central to both IBD and SpA pathogenesis. IL-23 is highly expressed in inflamed intestinal mucosa in IBD and synovial tissue in RA and SpA^35^ and drives Th17 differentiation and downstream production of IL-17, IL-6, TNF, and GM-CSF^36^. Macrophages and antigen-presenting cells are the principal sources of IL-23 in inflamed mucosa^37^, consistent with our observation that *E. faecalis* AHG0090 bioactive selectively suppresses IL-23 production in myeloid cells without broadly altering tissue immune cell composition.

At the molecular level, IL-23p19 expression downstream of toll-like receptor signalling is regulated in an NF-κB–dependent manner. Overexpression of IκB blocks IL-23p19 expression ^38^, and c-Rel has been identified as a critical transcriptional regulator of *Il23a*, with c-Rel-deficient dendritic cells failing to produce IL-23 in response to microbial stimuli^10^. Consistent with these studies, we demonstrate that *E. faecalis* AHG0090 bioactive suppresses c-Rel nuclear translocation, providing a mechanistic link between microbial bioactive activity and reduced IL-23 production. Importantly, this effect was conserved in human IBD patient-derived explants and macrophages, supporting translational relevance and validating patient explant systems as a powerful platform for mechanistic microbiome research.

Our findings highlight the limitations of compositional microbiome analyses and underscore the necessity of functional assays to define microbial contributions to disease. They further demonstrate that microbial ecology and growth conditions critically influence bioactive production, limiting the applicability of live bacterial therapeutics in immune-mediated disease. Instead, secreted microbial bioactive represent a tractable and mechanistically precise strategy to modulate upstream inflammatory pathways. In conclusion, we identify a microbiome-derived bioactive from *E. faecalis* AHG0090 that attenuates intestinal and joint inflammation through modulation of the IL-23–c-Rel NF-κB axis, supporting the development of bioactive-based therapies for gut and systemic inflammatory disorders.

## Methods

### Bacterial strains and culture conditions

*Enterococcus faecalis* AHG0090 was previously isolated from human faecal samples and characterised for its immunomodulatory properties. For preparation of cell-free culture supernatants (CS), *E. faecalis* AHG0090 was cultured under aerobic conditions at 37°C in brain heart infusion (BHI), M2GSC, or de Man, Rogosa and Sharpe (MRS) media as indicated in supplementary table 1. Cultures were grown to stationary phase, centrifuged to remove bacterial cells, and supernatants were filtered through 0.22 µm filters to ensure sterility. Filtered CS was stored at −80°C until use. BHI medium processed identically without bacterial growth served as control. For live bacterial administration, *E. faecalis* AHG0090 cultures were grown in BHI, washed, and resuspended in sterile phosphate-buffered saline (PBS) prior to gavage.

### Measurement of immunomodulatory activities

The LS174T-NF-kB*luc* reporter cell lines were used to test the immunomodulatory properties of bacterial CS and BHI media as previously described^34^. Briefly, LS174T reporter cells were stimulated treated with 10% v/v CS in complete DMEM medium prior with TNF stimulation (50 ng.ml^-1^) for 4 hours. NF-κB driven luciferase expression was assessed using the Pierce^TM^ Firefly Luc One-Step Glow Assay Kit (ThermoFisher Scientific) according to the manufacturer’s instructions.

### Animals

All animal experiments were approved by the University of Queensland Animal Ethics Committee 2022/AE000590 and conducted in accordance with national guidelines. *SKG* mice (ZAP-70 W163C mutant BALB/C) and *Winnie* mice were bred in-house under SPF conditions.

### SKG mouse model of ileitis and spondyloarthritis

12-16 week-old female SKG mice were gavaged with 1 x10^8^ CFU *E. faecalis* AHG0090 weekly for 4 weeks prior to curdlan (15mg/ml) injection (i.p). Disease activity including body weight loss and joint visual scores were recorded^39^. Briefly, the ankles were scored 0 for no swelling or redness, 1 for slight swelling and redness of digits, 2 for mild swelling and redness of footpad, joints and digits, 3 for severe swelling and redness of paw and digits and 4 for maximum swelling of paw, joints and digits. The maximum possible scores for visible inflammation are presented as the total of subscores up to 16.

### Treatment with E. faecalis AHG0090 supernatant in SKG SPF and Winnie mice

12-16 weeks old female SKG mice were challenged with curdlan intraperitoneally and subsequently treated with 200ul of CS from *E. faecalis* AHG0090 or BHI intraperitoneally biweekly for 2.5 weeks. Disease activity including body weight loss and visual joint scores were monitored as above.

Male and female Winnie mice were intrarectally gavaged with 50 µl of CS from *E. faecalis* AHG0090 and BHI for 14 days. Diarrhoea, rectal bleeding and body weight loss were monitored daily as previously described^34^. Diarrhoea scoring was interpreted as follows: 0 = no diarrhoea, solid stool; 0.5 = very mild diarrhoea, moist but formed stool; 1 = mild diarrhoea, formed but easily bisected by pressure applied; 1.5 = diarrhoea, no fully formed stools, and; 2 = severe, watery diarrhoea with minimal solid present.

### Histology

For histology scoring, the whole colon and ileum was rolled, fixed in 10% neutral buffered formalin, paraffin embedded, sectioned and stained with Haematoxylin and Eosin (H&E). Joints were fixed in 10% neutral buffered formalin and decalcified using EDTA prior to paraffin embedding, sectioning and H&E staining. Blind assessment of histologic inflammation (increased leukocyte infiltration, neutrophil counts, depletion of goblet cells, crypt abscesses, aberrant crypt architecture, increased crypt length, and epithelial cell damage and ulceration) for proximal, mid, and distal colon was performed as previously described.

### Gene expression

To quantify *in vivo* gene expression, the distal colon was snap frozen and homogenised in TRIzol. RNA was extracted using RNA extraction kits (Bioline) according to manufacturer’s instructions. RNA concentration was measured using a Nanodrop 1000 spectrophotometer and cDNA synthesis performed using 1 µg of RNA and the iScript cDNA synthesis kit (BioRad) according to the manufacturer’s instructions. The expression of genes of interest were analysed using quantitative real time PCR (qrt-PCR) as previously described^24,25^. C_t_ values were generated, and relative quantitation was determined by the ΔC_t_ method.

### Isolation of peritoneal exudate and lamina propria cells for fluorescent staining and sorting

For peritoneal exudate collection, mice were treated with 200ul of either BHI media or CS from AHG0090 for 24 hours before curdlan (0.5mg/mouse) was administered intra- peritoneally. Exudate was collected from peritoneum 18 hours post curdlan by flushing the peritoneal cavity with 5 ml of sterile PBS. Cells were centrifuged and stained for flow cytometry. For lamina propria prep, Winnie mice were treated with 50ul of either BHI or AHG0090 CS for 1hour. The colon was flushed and digested with EDTA (concentration, company) for 30mins followed by collagenase D (Roche) and DnaseI (Roche) digestion. Lamina propria cells were isolated using the Percoll (company) gradient centrifugation. Both exudate cells and lamina propria cells were stained with live/dead marker (zombie aqua) for live cell staining for 10mins, followed by surface markers from Biolegend : BV421 CD45.2 (Clone 104, 109832), BV605 CD11c (Clone N418, 117333), BV785 I-A/I-E (M5/114.15.2, 107645), FITC F4/80 (BM8, 123108,), PercpCy5.5 Ly6G (1A8, 127616),Pe-Cy7 CD11b (M1/70, 101216) and APC-Cy7 Ly6C(HK1.4, 128026) for 30mins. Cells were sorted using BD FACSAria Fusion Sorter. Sorted cells were lysed for RNA extractions using Qiagen RNAeasy mini kit.

### Patient samples

All patient samples were collected and processed as part of the Mater Inflammatory Bowel Disease Biobank in accordance with the Mater Health Services Human Research Ethics Committee (HREC 2016001782 & HREC/14/MHS/125)

### Peripheral Blood Mononuclear Cell (PBMC) isolation and immunomodulatory assays

PBMCs were isolated from UC and CD patients by Ficoll gradient density centrifugation as previously described. For the assays, 500,000 cells/ well were plated in a 96-well plate and treated with 10% v/v of CS from *E. faecalis* AHG0090 and BHI in RPMI medium for 30 minutes, followed by stimulation with rhTNFα (50 ng/ml). Monocyte-derived macrophages were generated by differentiation of CD14 monocytes and stimulated with lipopolysaccharide (LPS) in the presence or absence of AHG0090 CS. Cytokines were measured using the Legendplex inflammation panel (BioLegend) according to the manufacturer’s instructions.

### Patient derived explants

Colonic and ileal biopsies (3 x 3mm pinch biopsies) were collected from CD (n=8) and UC (n=8) patients. Biopsies were weighed and washed with PBS and cultured in complete RPMI (supplemented with 10% FBS, 1% glutamine and 1% penstrep) for 24 hours. The concentrations of IL-23, IL-6 and IL-8 in the supernatants were measured using ELISA (IL-6 and IL-8 (Biolegend) and IL-23 (R&D)) according to the manufacturer’s instructions. Cytokines levels were normalised per mg weight of the biopsies.

### Immunofluorescence and c-Rel localisation

Macrophages were stimulated with LPS in the presence of AHG0090 CS or BHI control. Cells were fixed, permeabilised, and stained for c-Rel (Cell signalling). Nuclear localisation of c-Rel was assessed by immunofluorescence microscopy and quantified using image analysis software Image J.

### Statistical analysis

Data are presented as mean ± SD. Statistical analyses were performed using appropriate parametric or non-parametric tests as indicated. Multiple comparisons were corrected where applicable. A *p* value <0.05 was considered statistically significant.

---

**Figure S1. Live *E. faecalis* AHG0090 does not improve gut and joint inflammation in the presence or absence of the gut microbiota A)** Changes in body weight **B)**Visual joint scores C-D) Blinded histological score of inflammation of the ileum and joints **E-F)** Representative H&E images of the ileum and joints. Each dots represent individual mouse. Figures are generated by pooling data from 2 independent experiments **G)** Effect of *E. faecalis* CS grown in different medium on TNFα stimulated LS174T NF-κB reporter cell (n=2 independent experiments)

**Fig S2. Gating strategy for colonic lamina propria A)** 12 weeks old Winnie mice were treated with 50ul BHI or CS from *E. faecalis* AHG0090 for 1 hour intra-rectally and the colon was used for lamina propria isolation. Cells were stained and population highlighted in red were sorted using flow cytometry. B) Pie -chart demonstrating the proportion of different cell types present in the lamina propria C) Bar-graph showing the proportion of each cell types present in both BHI and CS AHG0090 treated mice. D) sXBP-1 mRNA expression in sorted cells. Each dot represents individual mouse.

**Fig S3. Gating strategy for exudate from SKG mice A)** SKG mice were treated with 200ul of BHI and CS AHG0090 24 hours prior curdlan injection. Exudate was collected 18 hours post curdlan administration. Cells were stained and population highlighted in red were sorted using flow cytometry. B) Pie -chart demonstrating the proportion of different cell types present in the exudate C) Bar-graph showing the proportion of each cell types present in both BHI and *E. faecalis* AHG0090 CS treated mice. D) sXBP-1 mRNA expression in sorted cells. Each dot represents individual mouse.

**Figure S4. Patient demographics and cytokine profile.** A) PBMCs and explants from both UC and CD patients were utilised for explant studies. Demographics for patients are summarised. B) IL-8 secretion from the ileal and colonic explants. C)PBMCs from same patients were stimulated with LPS (10ug/ml) in the presence of BHI or *E. faecalis* AHG0090 CS for 24 hours and cytokines/chemokines secreted by PBMCs were normalised to cells only control and presented as log fold change.

## Author contributions

**Conceptualization:** R.G., J.B., **Methodology:** R.G., P.Ó C, A.S.B, **Investigation:** R.G., P.Ó C, A.S.B., **Data curation:** R.G, A.S.B., **Formal analysis:** R.G., **Resources:** M.M., R.T., J.B, **Supervision:** M.M., R.T., J.B., **Writing - original draft:** R.G., **Writing - review & editing:** R.G., A.S.B., P.Ó C., M.M., R.T., J.B.

