## Supplementary Fig for "Bioactive produced by *Enterococcus Faecalis* targets IL-23 signalling and protects against colitis and joint disease"

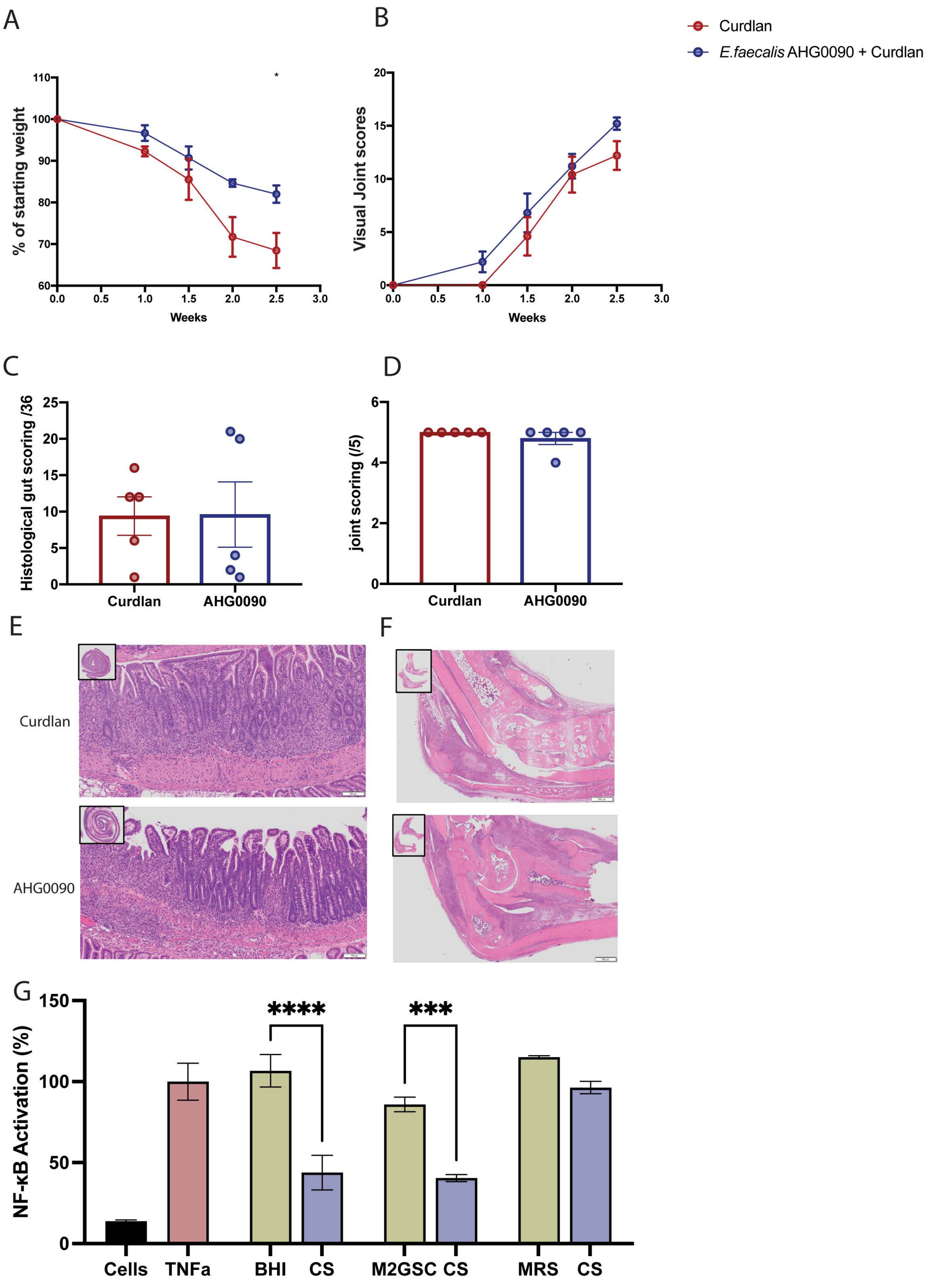
A
B
C
D
E
F
G
Curdlan
E.faecalis AHG0090 + Curdlan
% of starting weight
Weeks
Visual Joint scores
Histological gut scoring /36
joint scoring (/5)
Curdlan
AHG0090
NF-κB Activation (%)
****
***
Cells
TNFa
BHI
CS
M2GSC
MRS

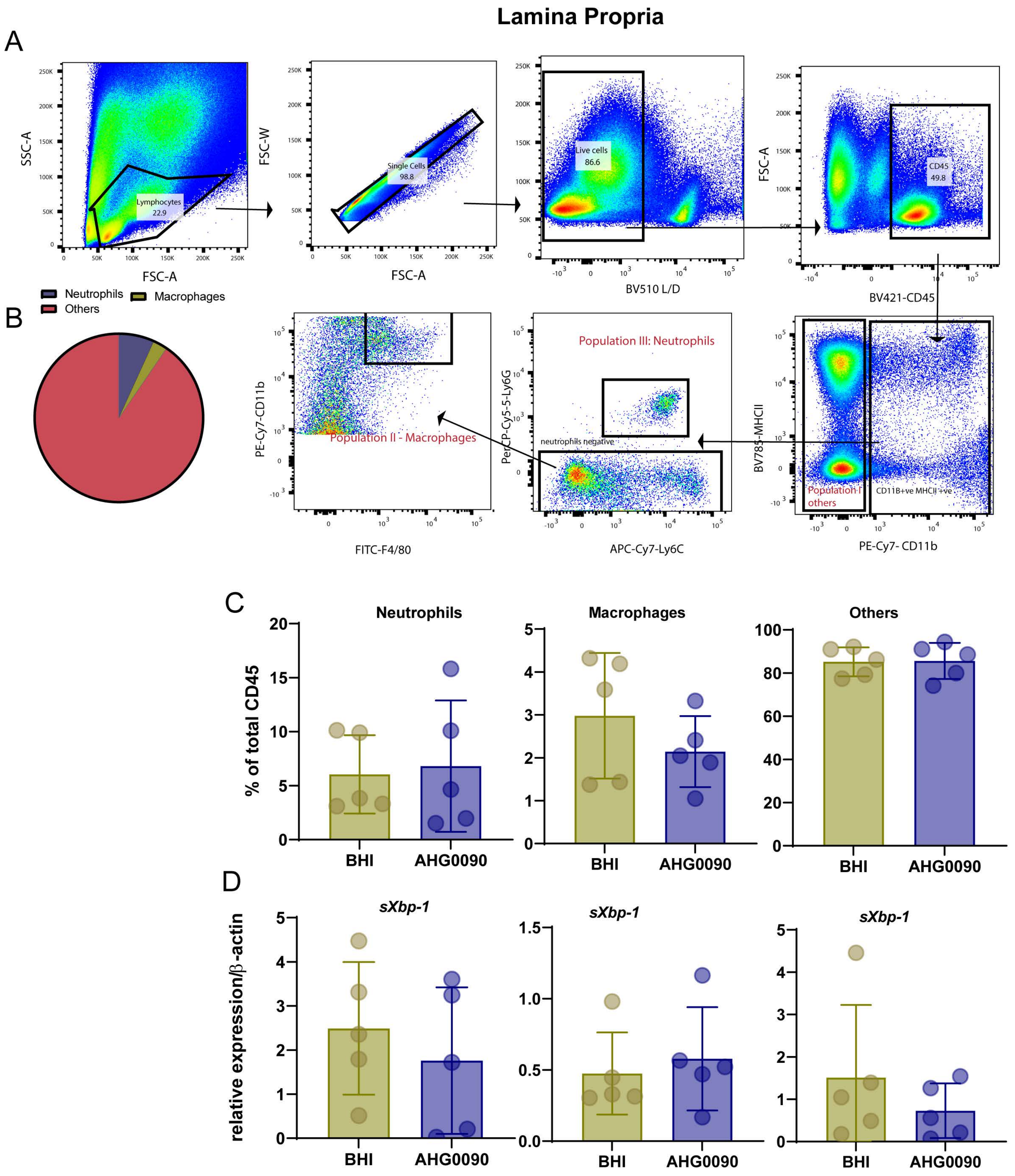

Lamina Propria
A
SSC-A
FSC-A
Lymphocytes
22.9
FSC-W
Single Cells
98.8
Live cells
86.6
BV510 L/D
CD45
49.8
BV421-CD45
B
Neutrophils
Macrophages
Others
PE-Cy7-CD11b
FITC-F4/80
Population II - Macrophages
PerCP-Cy5-5-Ly6G
APC-Cy7-Ly6C
Population III: Neutrophils
neutrophils negative
BV785-MHCII
PE-Cy7- CD11b
Population I
others
CD11B+ve MHCII+ve
C
% of total CD45
Neutrophils
Macrophages
Others
BHI
AHG0090
D
relative expression/β-actin
sXbp-1

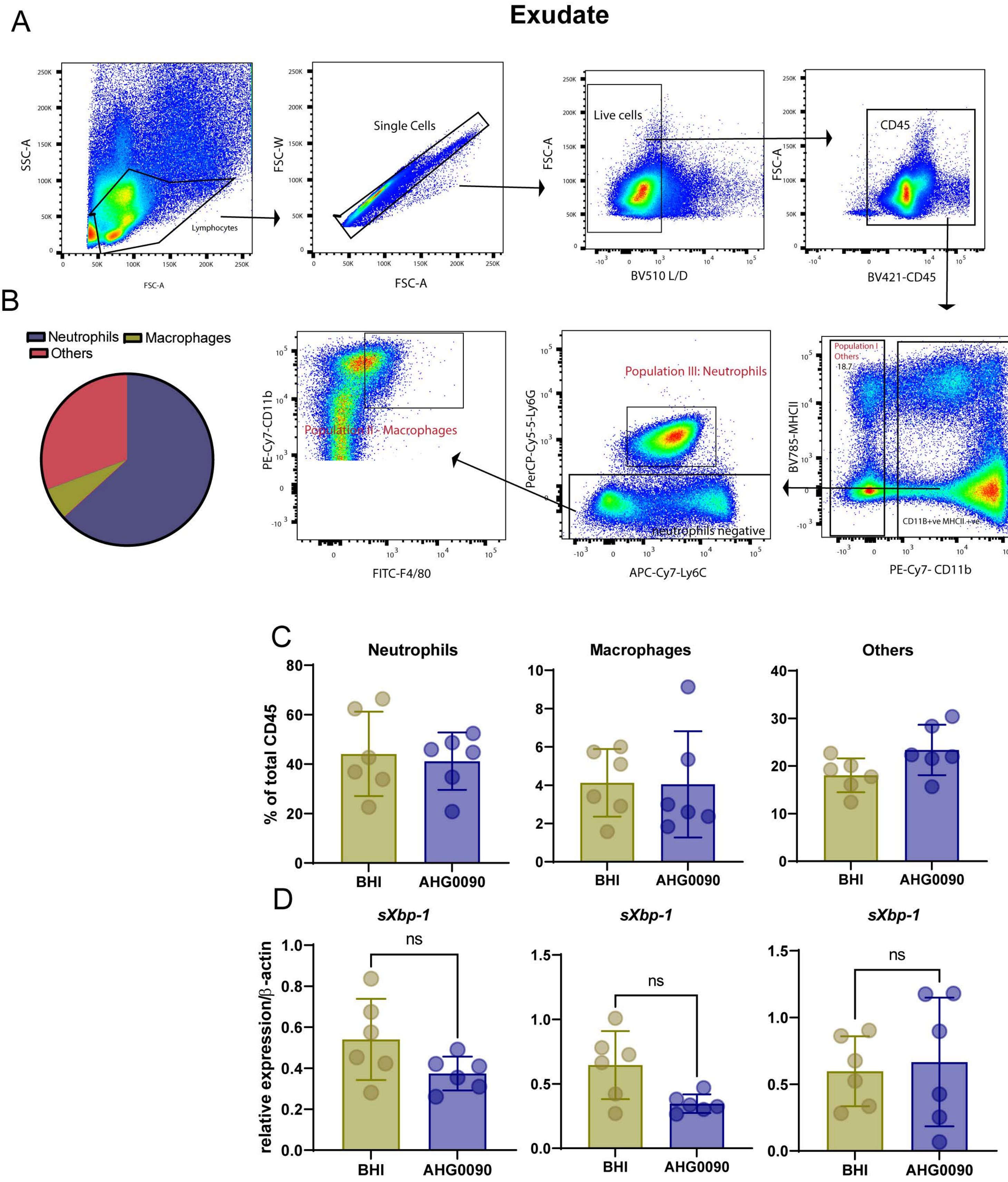
Exudate
A
SSC-A
FSC-A
Lymphocytes
FSC-W
Single Cells
FSC-A
Live cells
BV510 L/D
CD45
BV421-CD45
B
Neutrophils
Macrophages
Others
PE-Cy7-CD11b
Population II- Macrophages
FITC-F4/80
PerCP-Cy5-5-Ly6G
Population III: Neutrophils
neutrophils negative
APC-Cy7-Ly6C
BV785-MHCII
Population I
Others
18.7
CD11B+ve MHCII +ve
PE-Cy7- CD11b
C
Neutrophils
Macrophages
Others
% of total CD45
BHI
AHG0090
D
sXbp-1
ns
relative expression/β-actin

A

### Patient characteristics

| Characteristics | UC (n= 8 5F,3M) | CD (n=8 4M,4F) |
|---|---|---|
| **Age ± SD** | 51 ± 17 | 40 ± 16 |
| **Severity** | 4 none<br>2 mild<br>2 moderate | 3 none<br>3 mild<br>2 moderate |
| **Medication** | 5 IM, 5-ASA<br>2 none<br>1 Vedolizumab | 5 IM, anti-TNF<br>1 none<br>1 Ustekinumab |

B

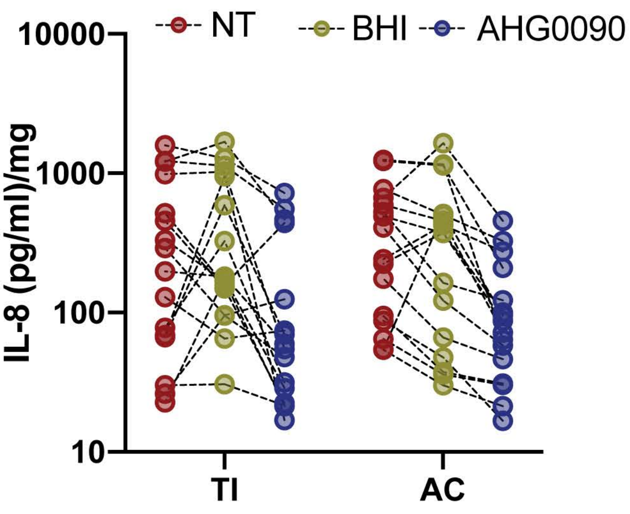


C

PBMCs

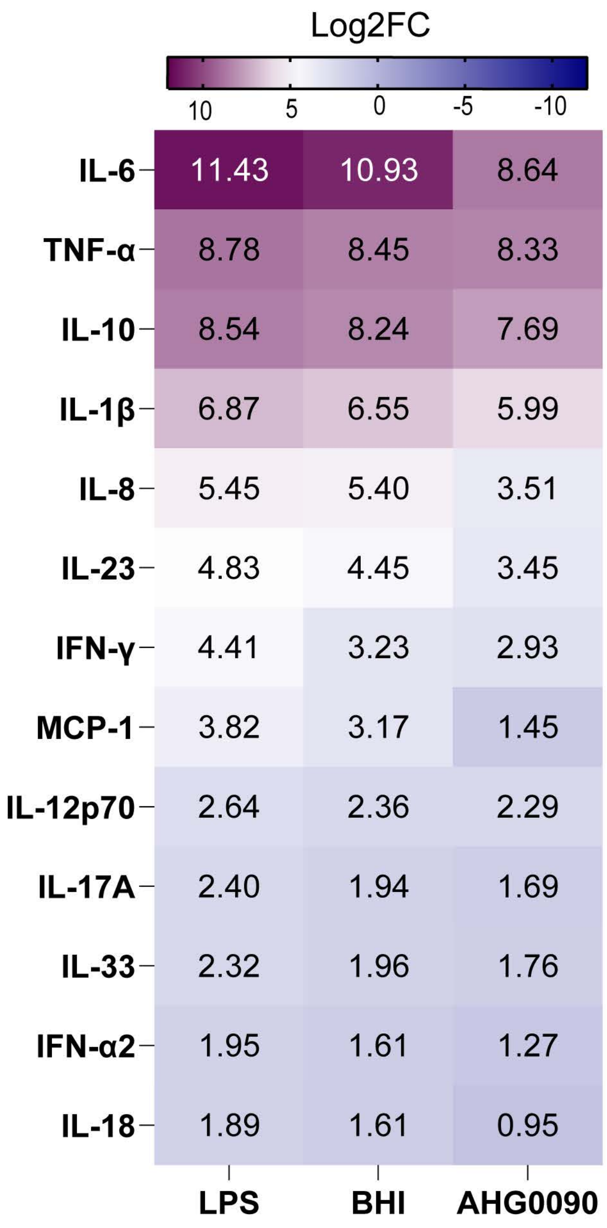
